# Dissection of the Hydrogen Metabolism of the Enterobacterium *Trabulsiella guamensis*: Identification of a Formate-Dependent and Essential Formate Hydrogenlyase Complex Exhibiting Phylogenetic Similarity to Complex I

**DOI:** 10.1101/563510

**Authors:** Ute Lindenstrauß, Constanze Pinske

## Abstract

*Trabulsiella guamensis* is a non-pathogenic enterobacterium that was isolated from a vacuum cleaner on the island of Guam. It has one H_2_-oxidizing Hyd-2-type hydrogenase (Hyd), and encodes a H_2_-evolving Hyd that is most similar to the uncharacterized *Escherichia coli* formate hydrogenlyase (FHL-2_*Ec*_) complex. The FHL-2_*Tg*_ complex is predicted to have 5 membrane-integral and between 4-5 cytoplasmic subunits. We could show that FHL-2_*Tg*_ complex catalyses the disproportionation of formate to CO_2_ and H_2_. FHL-2_*Tg*_ has an activity similar to the *E. coli* FHL-1_*Ec*_ complex in H_2_-evolution from formate, but the complex appears more labile upon cell lysis. Cloning of the entire 13 kbp FHL-2_*Tg*_ operon in the heterologous *E. coli* host has now enabled us to prove FHL-2_*Tg*_ activity unambiguously and allowed us to characterize the FHL-2_*Tg*_ complex biochemically. Although the formate dehydrogenase (FdhH) gene *fdhF* is not encoded in the operon, the FdhH is part of the complex and FHL-2_*Tg*_ activity was dependent on the presence of *E. coli* FdhH. Also, in contrast to *E. coli, T. guamensis* can ferment the alternative carbon source cellobiose, and we further investigated the participation of both the H_2_-oxidizing Hyd-2_*Tg*_ and the H_2_-forming FHL-2_*Tg*_ under these conditions.

**Importance:** Biological H_2_-production presents an attractive alternative for fossil fuels. But in order to compete with conventional H_2_-production methods, the process requires our understanding on the molecular level. FHL complexes are efficient H_2_-producers and the prototype FHL-1_*Ec*_ complex in *E. coli* is well studied. This paper presents the first biochemical characterisation of an FHL-2-type complex. The data presented here will enable us to solve the long-standing mystery of the FHL-2_*Ec*_ complex, allow a first biochemical characterisation of *T. guamensis*’s fermentative metabolism and establish this enterobacterium as model organism for FHL-dependent energy conservation.

## Introduction

Future use of molecular hydrogen (H_2_) as an energy carrier is a key step to replacing or reducing the anthropogenic use of fossil fuels. Sustainable H_2_ production from biological sources is one approach to tackle this problem. Many microorganisms produce H_2_ after fermentation of glucose but using glucose directly competes with human food production and requires the establishment of alternative carbon sources as substrates.

The apathogenic enterobacterium *Trabulsiella guamensis* is able to grow on both cellobiose and glucose, fermenting these through the mixed acid fermentation (1). *T. guamensis* was isolated from the dust of a vacuum cleaner in a library on Guam/USA (1). Phylogenetically related strains have also been isolated among cellulolytic microbes (2). It is phylogenetically most closely related to the pathogenic enterobacterium *Salmonella enterica*. Its genome sequence has been made available within the context of the project ‘Assembling the Tree of Life’ to increase the breadth of genome sequence data available for the enterobacteria (https://atol.genetics.wisc.edu/index.php). With regard to its [NiFe]-hydrogenase (Hyd) complement, the genomic data predict that the organism encodes only one uptake Hyd and one enzyme complex similar to the formate hydrogenlyase 2 (FHL-2_*Ec*_) complex of *Escherichia coli*.

The FHL-2_*Ec*_ complex is related to the H_2_-producing FHL-1_*Ec*_ complex, which catalyzes the oxidation of formate to CO_2_ and H_2_. The FHL-1_*Ec*_ complex is the main hydrogen-producing enzyme complex in *E. coli* under fermentative growth conditions and detoxifies formate. It consists of a formate dehydrogenase (FdhH) for formate oxidation and three iron sulfur-cluster-containing proteins, HycB, HycF and HycG, that transfer electrons to the hydrogenase subunit (HycE). It also has two membrane subunits, HycC and HycD, that are not involved in the redox reaction, but are essential for full FHL-1_*Ec*_ activity (Fig. 1) (3). The genes coding for the hydrogenase subunits and protein specific chaperones of FHL-1_*Ec*_ are in the *hycA-I* operon. The FdhH protein is encoded by the *fdhF* gene, but its location on the chromosome is not in the vicinity of the *hyc* genes. The FHL-1_*Ec*_ architecture resembles the proton-translocating Complex I of the respiratory chain (4, 5). Unfortunately, our previous attempts to prove the predicted proton translocation ability of the FHL-1_*Ec*_ complex have remained unfruitful (6).

**Figure 1:**
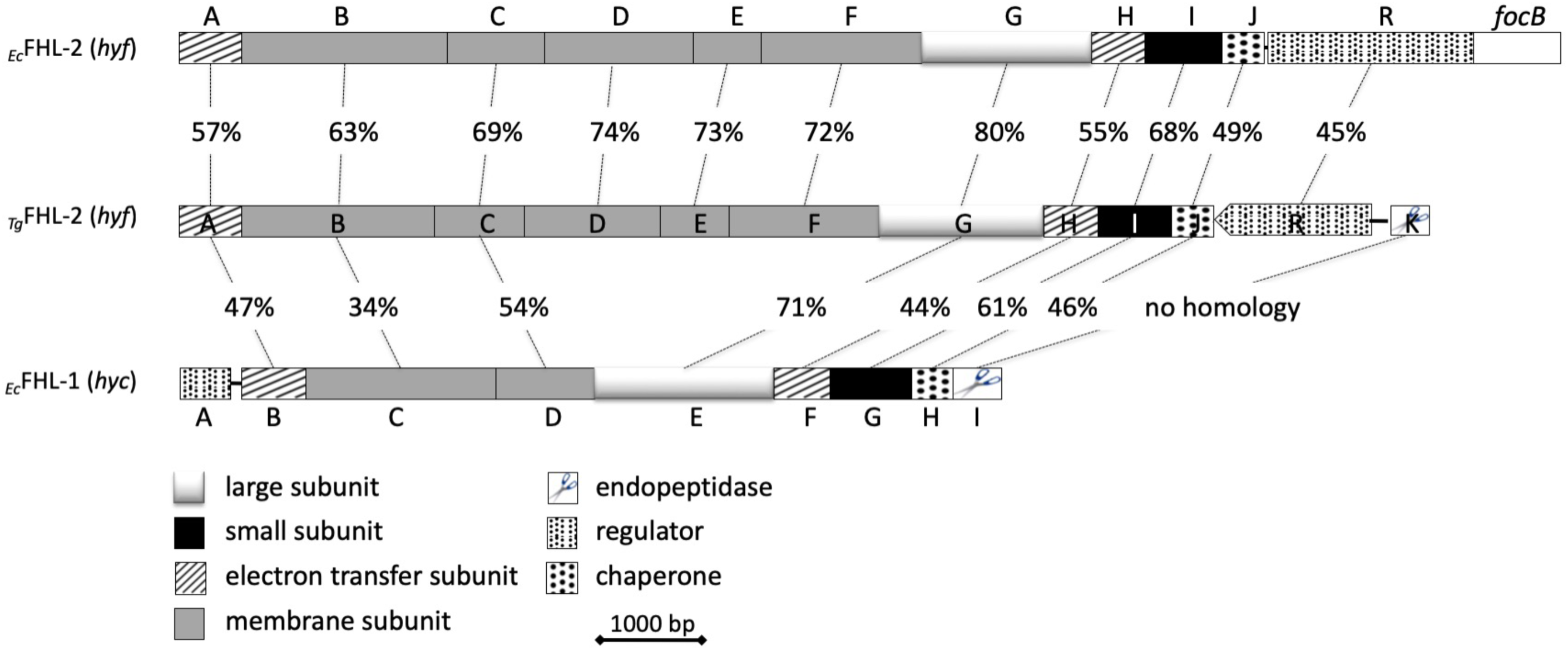
Operon structure of formate hydrogenlyase complex-encoding genes from *E. coli* and *T*. *guamensis*. The numbers indicate the percentage of identical amino acid residues shared between the corresponding proteins.

The second FHL complex (FHL-2_*Ec*_) is predicted to have five instead of two membrane subunits. The FHL-2_*Ec*_ complex is encoded by the *hyfA-R* genes, but neither gene expression nor protein activity for this FHL-2_*Ec*_ complex was yet directly demonstrated (7–11). It was further predicted that this complex might also rely on FdhH as electron donor (8). The two FHL complexes can be unambiguously distinguished based on the presence of the FHL-2-specific protein HyfE (Fig. 1).

The *T. guamensis* FHL-2_*Tg*_ complex encoded by the *hyf*-operon is also predicted to have five membrane subunits and also no formate dehydrogenase is encoded within the vicinity of the *hyf* operon, but is encoded elsewhere on the chromosome. Notably, however, the FHL-2_*Tg*_hydrogenase large subunit phylogenetically clusters together with the corresponding *E. coli* complex I protein NuoCD and its FHL-2_*Ec*_ catalytic subunit HyfG (Supp. Fig. S1)(12). Other aspects of the *T. guamensis hyf*-operon composition are highly similar to the *hyf* counterpart in *E. coli* but with significant variation in the organisation of associated transcriptional regulators, protein maturation factors and the surrounding genome context (see Fig. 1). For example it encodes an endoprotease gene *hyfK*, whose gene product is predicted to be required for the processing of the large subunit after completion of the metal-cofactor insertion and is missing in the *E. coli hyf*-operon (13). Instead, there is no formate transporter FocB encoded within the *T. guamensis hyf*-operon. The *E. coli* FocB protein was presumed to participate in formate transport, but the main formate transporter in *E. coli* is FocA, located within the *focA-pflB* operon, encoding also the pyruvate formate-lyase (14–16). The *focA-pflB* operon is also conserved in *T. guamensis* and the FocA protein sequence is 92% identical to FocA in *E. coli*. Taken together, these features suggest an apparent functionality of the FHL-2_*Tg*_ complex.

The number and complement of the H_2_-oxidizing Hyd vary significantly among enterobacteria. The *E. coli* Hyd-1 and Hyd-2 both catalyse H_2_-oxidation and face the periplasmic side of the cytoplasmic membrane (5, 17). Although their respective catalytic subunits HyaB and HybC share 40% identity at the amino acid level, the architecture of the entire membrane complex differs significantly, conferring upon them unique enzymic properties. The Hyd-1 complex has three different subunits and can function in the presence of oxygen (18). Hyd-2 on the other hand combines four different subunits (HybOABC) and is highly catalytically active (18). Hyd-2 is also involved in the generation of a proton gradient during catalysis (19).

*S. enterica* has 3 H_2_-oxidizing hydrogenases (catalytic subunits HyaB, HybC and HydB in Supp. Fig. S1) and one FHL-1 complex (catalytic subunit HycE in Supp. Fig. S1), and synthesis of these enzymes varies depending on its environmental situation (20). For example, in *S. enterica* and the pathogenic epsilon-proteobacterium *Helicobacter pylori* energy-conserving H_2_-oxidation contributes to the pathogenicity of both bacteria, because anaerobic respiration enhances survival of the pathogen in the host (21–23). Interestingly, *T. guamensis* encodes only one H_2_-oxidizing hydrogenase, which is similar to Hyd-2 of *E. coli* (HybC in Supp. Fig. S1)(1).

The *T. guamensis* hydrogenase complement now poses the unique opportunity to study H_2_ production by an organism than only has the FHL-2_*Tg*_ complex. Thus, we can gain considerable insight into the bioenergetics of this intriguing complex and we can determine what deficiencies in the FHL-2_*Ec*_ complex prevent it being active under standard laboratory conditions.

## Results and discussion

### Growth conditions that favour Hyd-2_*Tg*_ activity

*T. guamensis* can utilize cellobiose, the disaccharide derived from cellulose, for anaerobic growth with a growth rate of μ = 0.20 h^-1^ (Fig. 2A). The growth rate with glucose was more than twice that of the cellobiose growth with μ = 0.57 h^-1^. The growth rate of *E. coli* with glucose was μ = 0.24 h^-1^, which is in the same order of magnitude as the cellobiose growth rate of *T. guamensis*, while no growth of *E. coli* was observed with cellobiose. *T. guamensis* encodes a β-D-glucosidase similar to BglC from *Bacillus subtilis*, which is not present in *E. coli* and a periplasmic β-D-glucosidase BglX whose function has not been determined in *E. coli*, giving it the capacity to use cellobiose. Extracts from these cultures were applied to native-PAGE and stained for Hyd activity, which revealed a H_2_-oxidation activity as staining of a red band. In this assay, *E. coli* showed a fast-migrating Hyd-1 activity band, two slower migrating bands that can be assigned to Hyd-2 and one band belonging to the FHL-1_*Ec*_ complex (24) (Fig. 2B). Weaker bands can be observed near the top of the gel, which were previously identified as due to weak H_2_ oxidation activity catalysed by formate dehydrogenases N and O (25).

**Figure 2:**
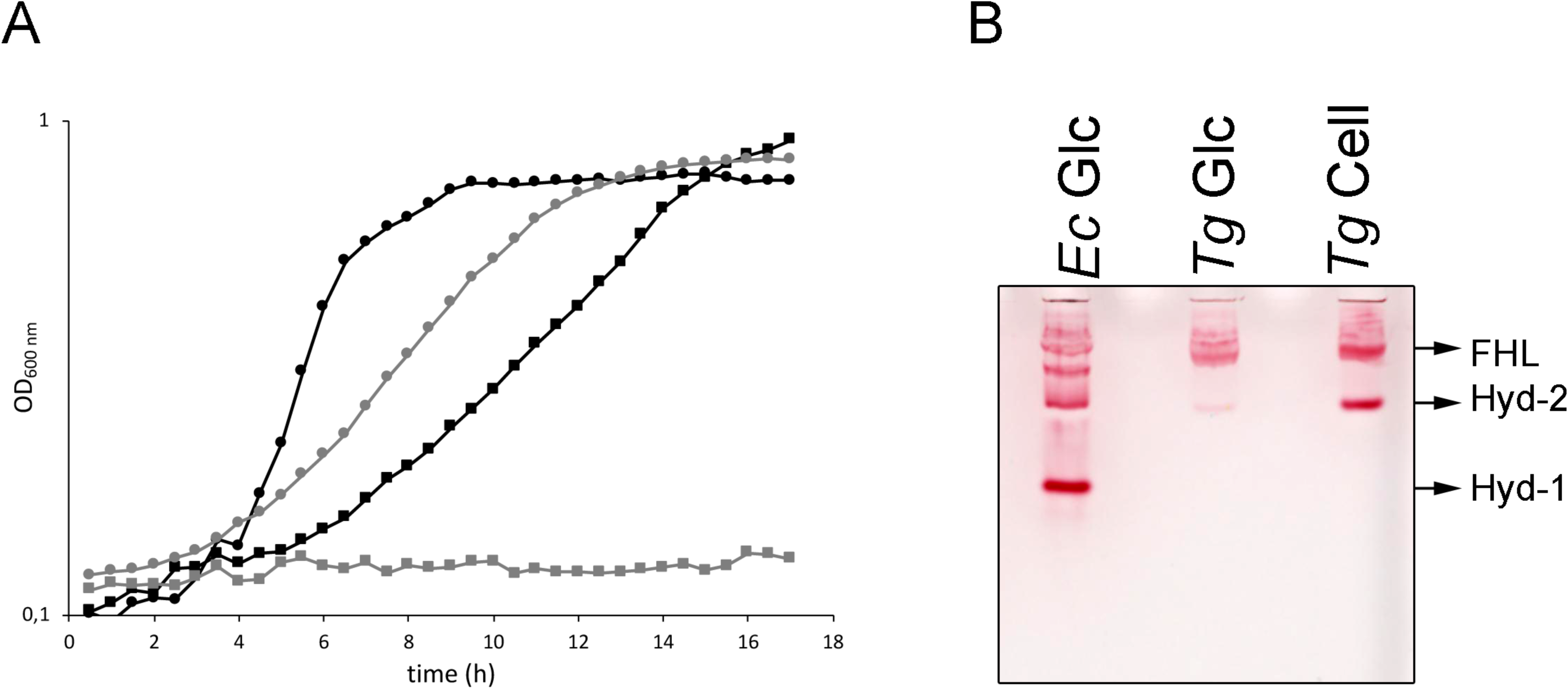
Growth and hydrogenase enzyme synthesis during cellobiose fermentation. (A) Cells of *T. guamensis* (*Tg* in black) and *E. coli* strain MG1655 (*Ec* in grey) were grown as triplicates in standing liquid cultures in M9 minimal medium with either 0.8% (w/v) glucose (circles) or cellobiose (squares) as sole carbon and energy source. Growth was monitored in a Tecan plate-reader. (B) Cells were harvested at the beginning of stationary phase and 25 μg protein separated on a 7.5% native-PAGE and subsequently stained for hydrogenase activity with benzyl viologen, 2,3,5-triphenyltetrazolium chloride and H_2_. The active hydrogenase complexes are labelled on the right of the gel based on the migration pattern of the *Ec* enzymes.

The *T. guamensis* extracts show an intensively staining red band under both growth conditions and this migrates at approximately the same position as the FHL-1_*Ec*_ complex. Furthermore, a faster-migrating band is weakly visible in glucose-grown cells but strongly increased in intensity in cellobiose-grown cells. Based on the homology to the *E. coli* Hyds, the lower band likely represents Hyd-2. This apparent glucose repression of Hyd-2 is rather surprising, because synthesis of the *E. coli* Hyd-2 is not repressed during mixed acid fermentation as sufficient fumarate is produced internally that can be reduced to succinate with electrons derived from H_2_ (26).

In order to unambiguously verify the identity of these bands, a Hyd-2-deficient strain was constructed that carried the insertion of a kanamycin-resistance cassette within HybC, the catalytic subunit of Hyd-2. Strain TGH001 and its parental strain were grown in glucose, cellobiose and glycerol/fumarate medium and extracts applied to a native-PAGE (Fig. 3A). This experiment verified that the faster migrating band is indeed Hyd-2_*Tg*_. It additionally confirmed that the activity of the enzyme is higher after growth with cellobiose and especially high in the presence of glycerol and fumarate. Furthermore, the Hyd-2_*Tg*_ coding genes were introduced into an *E. coli* strain (DLH04) that lacked the genes for the structural subunits of Hyd-2 *hybOABC* (Fig. 3B). In addition, this strain is devoid of Hyd-1 and FHL-1_*Ec*_ activity. When introduced, the plasmid p*Tg*-hybOG resulted in an activity band that migrated slightly slower than that of the parental enzyme. This retardation is presumably caused by the addition of the histidine affinity tag on HybC and has been previously observed for other Hyds in native-PAGE (27). The plasmid carries the enzyme-specific maturation enzymes, but lacks the genes for the general maturation machinery of the [NiFe]-cofactor, indicating that the *E. coli* maturation machinery recognizes the foreign HybC protein and introduced the [NiFe]-cofactor. Induction of gene expression by IPTG resulted in the absence of the activity band, indicating that high amounts of protein have detrimental effects, possibly caused by an overburdening of the maturation machinery or the membrane-targeting apparatus.

**Figure 3.**
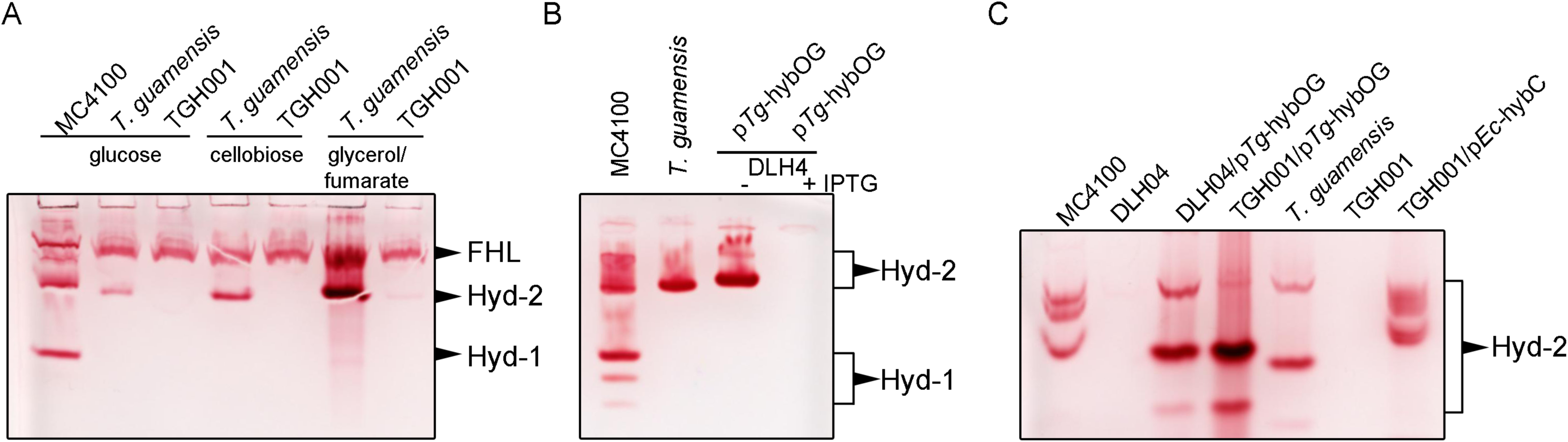
Identification of _*Tg*_Hyd-2: Cells of strains MC4100, *T. guamensis* and TGH001 (*Tg hybC*∷*kan*) were grown anaerobically in (A) M9-medium with either 0.8% (w/v) glucose, 0.4% (w/v) cellobiose or 0.4% (v/v) glycerol and 15 mM fumarate or (B)/(C) TYEP with 0.4% (v/v) glycerol and 15 mM fumarate, as indicated. Panel (B) additionally shows *E. coli* strain DLH04 (MC4100 DE3 Δ*hycA-I* Δ*hybO-C* Δ*hyaB*) and after complementation with p*Tg*-hybOG plasmid. The + and – indicate addition or absence of 0.2 mM IPTG during growth. Panel (C) additionally shows strain TGH001 (*T. guamensis hybC*∷*kan*) complemented with plasmids p*Tg*-hybOG and p*Ec*-hybC. In this panel all bands represent Hyd-2 activities. Extracts of 25 μg protein were applied to native-PAGE and subsequently stained for Hyd-activity. The migration pattern of the Hyd is indicated on the right.

Based on the findings above, the TGH001 (*Tg hybC*∷*kan*) strain was used to test if the introduced mutation was the cause for the phenotype. Therefore, the growth conditions with glycerol and fumarate were chosen to favour optimal Hyd-2 synthesis and also the gel composition was slightly adjusted, so that it revealed a second, slower migrating band for Hyd-2 in *T. guamensis* (Fig. 3C, lane 5), which was absent in TGH001 (Fig. 3C, lane 6) and even three bands belonging to Hyd-2 in *E. coli* (lane 1). These multiple bands represent various subcomplexes of Hyd-2 that differ in the presence or absence of the membrane anchor subunits HybA and HybB (28). A further unspecific faster migrating band of Hyd-2 can be observed on the bottom of the gel after over-expression and might represent a form of degraded Hyd-2, similar to that observed for Hyd-1 when over-expressed (29). For the complementation, strain TGH001 was transformed either with plasmid p*Tg*-hybOG or with p*Ec*-hybC. Analysis on native-PAGE showed that the main activity band of the p*Tg*-hybOG construct returned in the complemented strain TGH001 (Fig. 3C, lane 4), but the migration pattern was slightly slower than that of the untagged complex in *T. guamensis* (Fig. 3C, lane 5) but identical to the activity band seen of the His-tagged version in *E. coli* strain DLH04 (Fig. 3C, lane 3). The *E. coli hybC* gene from plasmid p*Ec*-hybC can also complement the lack of Hyd-2 activity in strain TGH001 (Fig. 3C, lane 7). Surprisingly, the activity band pattern of the chimeric complex in *T. guamensis* resembled that of *E. coli* Hyd-2 (Fig. 3C, lane 1) with the exception of the slower migration due to the fused StrepII-affinity tag on HybC. This finding would imply, that the large subunit co-ordinates assembly of the entire tetrameric complex, although in an alignment the *Tg*HybC and *Ec*HybC proteins have only 15 amino acid positions that are entirely different and those mostly locate on the solvent exposed outer surface, except for positions 132 and 145 that face the respective other HybC protein. More research with purified chimeric complexes will be necessary to resolve this question, but currently the affinity tags are not suitable for isolation of the entire Hyd-2 complex.

### Hyd-2_*Tg*_ is a H_2_-oxidizing Hyd

The cultures from the different growth media were tested for their H_2_-production, determined as H_2_ accumulation in the headspace above the culture (Tab. 1). When cells were grown with glucose, they produced the highest amount of H_2_ with 48 μmol OD_600 nm_^-1^ ml^-1^. This is about 60% of that produced by an *E. coli* culture under the same conditions but in the same range as that of strain TGH001 (*hybC*∷*kan*) (Tab. 1). The latter result indicates that Hyd-2 does not contribute to H_2_ production under these conditions and is only responsible for a small fraction of re-oxidation of the product. The amount of H_2_ was reduced to 17% when *T. guamensis* was grown in the presence of cellobiose, and was absent when the cells were grown with glycerol/fumarate. Here the extent of the re-oxidation of H_2_ by Hyd-2 becomes evident, because strain TGH001 (*hybC*∷*kan*) produced more than twice the amount of H_2_ than the wild type grown with cellobiose and generated even higher amounts in glycerol/fumarate medium.

**Table 1:** H_2_-evolution from cultures

| | $\mu\text{mol H}_2 \text{ OD}_{600 \text{ nm}^{-1} \text{ ml}^{-1}}$ | | | |
| --- | --- | --- | --- | --- |
|  | *T. guamensis* | TGH001 | TGH001/p*Tg*-hybOG | MC4100 |
|  |  | *(hybC::kan)* |  |  |
| Glucose | 48.3 ± 9.1 | 51.5 ± 2.0 | 31.9 ± 2.3 | 82.8 ± 2.7 |
| Cellobiose | 8.2 ± 3.9 | 19.9 ± 5.4 | <0.1 | n.a. |
| Glycerol/fumarate | <0.1 | 29.9 ± 2.3 | <0.1 | <0.1 |

n.a. – not available

### Formate is the substrate for the labile FHL-2_*Tg*_ complex

Cell suspensions of *T. guamensis* and *E. coli* strain MC4100 were applied to a Clark-type electrode and formate was tested to initiate H_2_-production. The activity of the cells was in a similar range as that of *E. coli* with 3.21 ± 0.24 and 3.08 ± 0.79 μmol min^-1^ mg protein^-1^, respectively (Tab. 2). The mutant TGH001 (*hybC*∷*kan*) with the insertion that inactivated the Hyd-2 enzyme, showed slightly reduced H_2_-production with 2.61 ± 0.24 μmol min^-1^ mg protein^-1^. This clearly demonstrated that the remaining activity was due to FHL-2_*Tg*_.

**Table 2:**
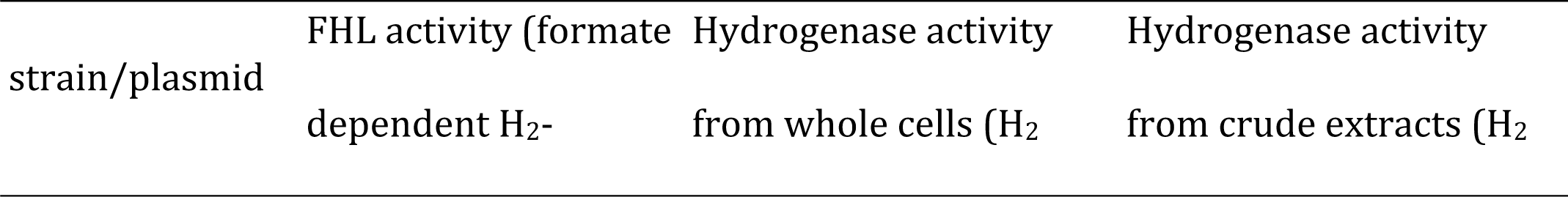
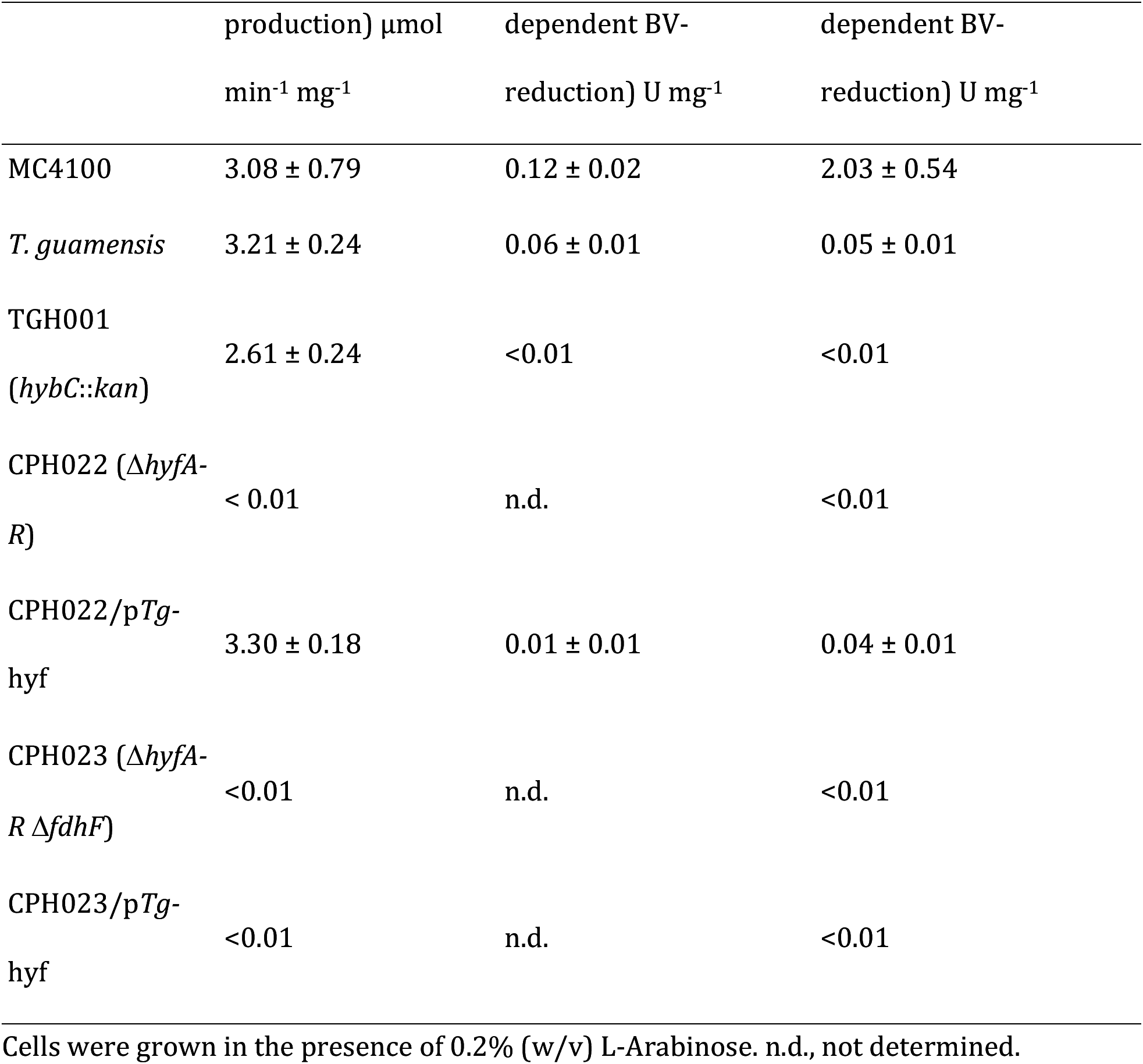
Hydrogenase activities from FHL-2_*Tg*_

| strain/plasmid | FHL activity (formate dependent H<sub>2</sub>- | Hydrogenase activity from whole cells (H<sub>2</sub> | Hydrogenase activity from crude extracts (H<sub>2</sub> |
| --- | --- | --- | --- |

| | production) $\mu\text{mol}$<br>$\text{min}^{-1} \text{mg}^{-1}$ | dependent BV-reduction) $\text{U mg}^{-1}$ | dependent BV-reduction) $\text{U mg}^{-1}$ |
| --- | --- | --- | --- |
| MC4100 | $3.08 \pm 0.79$ | $0.12 \pm 0.02$ | $2.03 \pm 0.54$ |
| *T. guamensis* | $3.21 \pm 0.24$ | $0.06 \pm 0.01$ | $0.05 \pm 0.01$ |
| TGH001<br>(*hybC::kan*) | $2.61 \pm 0.24$ | <0.01 | <0.01 |
| CPH022 ($\Delta\text{hyfA-}$<br>*R*) | < 0.01 | n.d. | <0.01 |
| CPH022/p*Tg-*<br>*hyf* | $3.30 \pm 0.18$ | $0.01 \pm 0.01$ | $0.04 \pm 0.01$ |
| CPH023 ($\Delta\text{hyfA-}$<br>*R* $\Delta\text{fdhF}$) | <0.01 | n.d. | <0.01 |
| CPH023/p*Tg-*<br>*hyf* | <0.01 | n.d. | <0.01 |

Cells were grown in the presence of 0.2% (w/v) L-Arabinose. n.d., not determined.

Experiments showing the optimum pH for activity revealed that when cells were grown at an initial pH of 5.5, the activity was up to 220% higher than when the medium pH was initially set to 6.5 or 7.5 (Supp. Tab. 1). However, the assay conditions showed that of the three pHs tested, the highest activity was obtained when cells were assayed at pH 6.5. These data clearly show that the activity is most stable under slightly acidic pH, but overall enzyme activity is highest when the cells try to counteract acidification and formate accumulation during growth, a function that was originally proposed for FHL-1_*Ec*_ (30).

Using *E. coli* as chassis, the cloned *T. guamensis hyfA-K* operon was introduced into *E. coli* strain CPH022, which is devoid of the entire operons encoding FHL-1_*Ec*_ and FHL-2_*Ec*_. The *T. guamensis hyfA-K* operon showed H_2_ production from the pBAD24 backbone, and this was entirely dependent on the presence of L-Arabinose (Supp. Fig. 2). This plasmid restored an FHL activity on the electrode to 3.30 ± 0.18 μmol min^-1^ mg protein^-1^ to strain CPH022 (Tab. 2). Furthermore, deletion of the *fdhF* gene, coding for the FHL-1_*Ec*_ associated formate dehydrogenase H, and subsequent addition of p*Tg*-hyf showed that the activity of the heterologous FHL-2_*Tg*_ complex was absent and thus identified FdhH as the electron input module for FHL-2_*Tg*_ in *E. coli*. The *E. coli* and *T. guamensis* formate dehydrogenases are both encoded at distinct locations from the operons encoding FHL and recent work indicated that FdhH can form several protein complexes within the cell (31). The proteins share 95% amino acid identity between the two organisms, including the catalytic selenocysteine residue. This conservation is considerably higher than that of the remaining components of the complexes (Fig. 1). The *E. coli* FdhH can therefore substitute for the *T. guamensis* FdhH and provides electrons for H_2_-production from formate.

The extracts from the heterologous expression of the FHL-2_*Tg*_ complex were further tested in a spectrophotometric assay, which measure total hydrogenase activity of all Hyd, including FHL-1_*Ec*_, as H_2_-dependent reduction of the artificial electron dye benzyl viologen (27). Performing the assay with whole cells allowed quantification of the periplasmically oriented Hyd enzymes like Hyd-1 and Hyd-2 only, because oxidized BV does not diffuse across the intact membrane (28). This assay resulted in activities of 0.12 and 0.06 U mg protein^-1^ for *E. coli* and *T. guamensis*, respectively (Tab. 2). Cell lysis increased the activity of the *E. coli* extract to 2.03 U mg protein^-1^ due to the accessibility of the FHL-1_*Ec*_ complex. This increase was neither detectable in extracts of *T. guamensis* nor in extracts of CPH022 synthesizing FHL-2_*Tg*_ heterologously (Tab. 2). This result either implies that the FHL-2_*Tg*_ complex is unstable upon cell lysis or unable to perform the reverse H_2_-oxidation reaction in the presence of high amounts of H_2_.

The strain TGH002 carries a deletion of the entire *hyfA-K* operon and could only be stably obtained as a merodiploid in the presence of an *in trans* copy of the same operon. The regulation of the plasmid-encoded operon was under the control of an arabinose-inducible promoter. Numerous attempts to generate the mutant without this extra copy failed. Similarly, anaerobic growth of TGH002 without arabinose was not possible. Low amounts of L-Ara (0.05% w/v) were already sufficient to induce H_2_-production, but an increase of up to 2% (w/v) had no further significant effect on the H_2_-production showing that the amount of active FHL-2_*Tg*_ complex cannot be further increased (Fig. 4). This is similar to the observation of Hyd-2 activity, which cannot be increased when induced with IPTG. In contrast, the same plasmid showed that in *E. coli* strain CPH022 absence of L-Ara still showed low amounts of H_2_ (Fig. 4), rendering this construct unsuitable for further experiments in *T. guamensis*. In strain CPH022, the optimal L-Ara concentration for H_2_ production was between 0.05 and 0.1% (w/v), with a further increase of L-Ara causing a decrease in H_2_ accumulation. Similar to Hyd-2, over-production left the protein mainly inactive.

**Figure 4:**
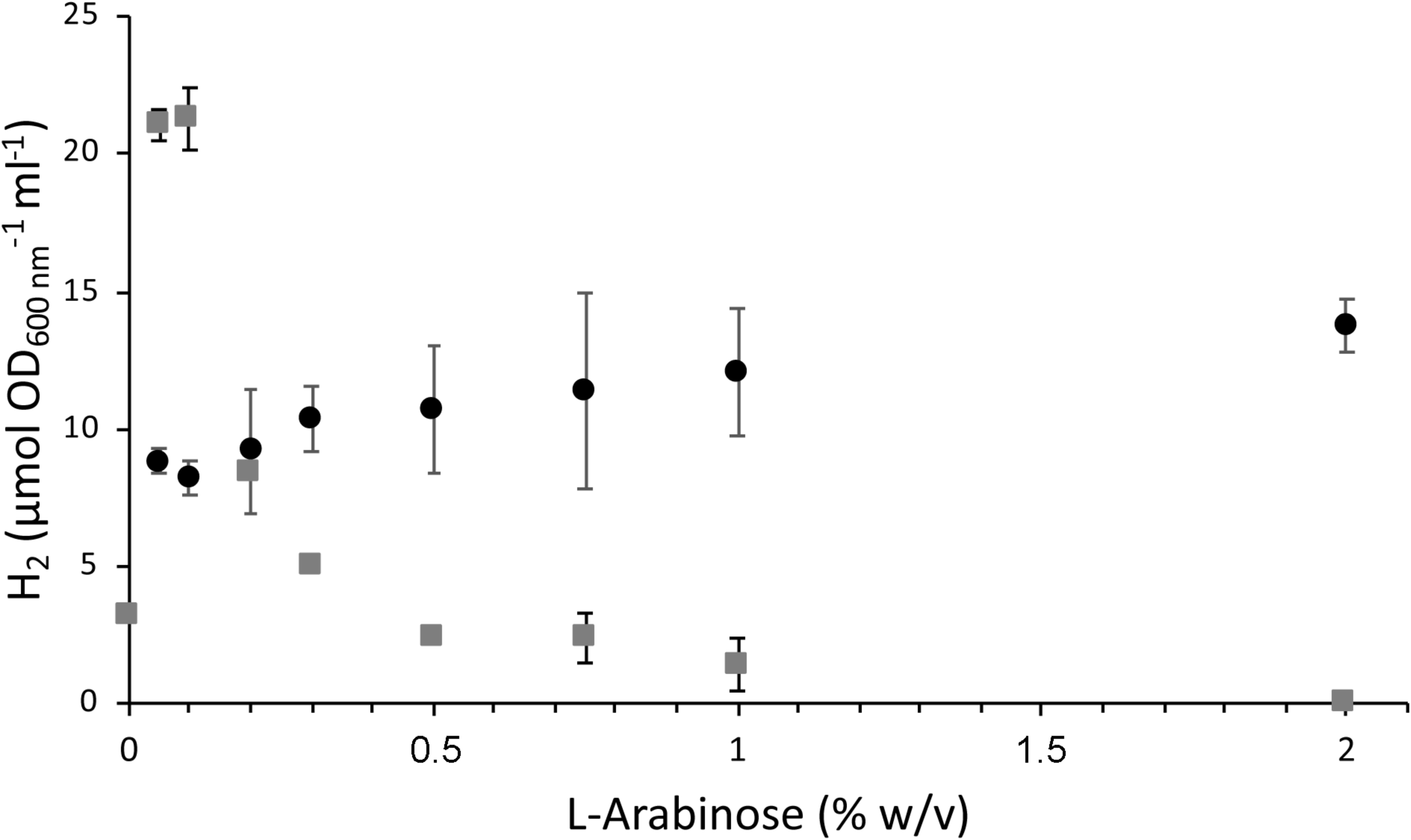
L-Arabinose-dependent H_2_-production from FHL-2_*Tg*_ in TGH002 (black circles) and CPH022 (grey squares). Cells were grown as liquid cultures in TGYEP medium, pH 6.5 in triplicates with the indicated amount of L-Ara in the growth medium. Headspace H_2_ content was determined by gas chromatography as described in the Materials and methods section.

## Conclusions

We show in this study that the FHL-2_*Tg*_ of the enterobacterium *T. guamensis* functions as a highly efficient H_2_-producing enzyme complex using formate as substrate. The activity is similar to that of FHL-1_*Ec*_ and it is dependent on the formate dehydrogenase H protein in *E. coli*. The number of subunits in the soluble domain between these two complexes is the same, but the difference is that five membrane subunits are present in the FHL-2_*Tg*_ complex, rather than the two in FHL-1_*Ec*_. It appears from the lack of growth of the *T. guamensis* Δ*hyfA-K* mutant without concomitant co-expression of the operon *in trans* that the FHL-2_*Tg*_ complex is essential in *T. guamensis*. This could suggest that the extra membrane subunits HyfDEF are required for generation of a proton gradient in *T. guamensis*.

The Hyd proteins can be functionally interchanged between *E. coli* and *T. guamensis*, and this provides an excellent model system to investigate the proton-translocating potential of the FHL-2_*Tg*_ in *E. coli* where it is not essential. Moreover, because our experiments show that *T. guamensis* is genetically tractable, this bacterium can be then used to re-introduce genetically modified FHL-2_*Tg*_ complexes to determine what makes FHL-2_*Tg*_ essential.

## Materials and methods

### Medium and growth conditions

*T. guamensis* strain DSM16940 was obtained from the DSMZ, Germany. *E. coli* strain MC4100 was used for experiments throughout this study (32). The strain DLH04 (MC4100 DE3 Δ*hycA-I* Δ*hybO-C* Δ*hyaB*) was described (D. Lubek, A. H. Simon and C. Pinske, submitted for publication) (Tab. 3). Strains of *E. coli* and *T. guamensis* were routinely grown in liquid LB medium (33) solidified with 1.5% (w/v) agar for plate growth. Antibiotics were added to a final concentration of 100 μg ml^-1^ for ampicillin, 30 μg ml^-1^ for chloramphenicol and 50 μg ml^-1^ for kanamycin. Unless otherwise stated, L-arabinose was added to a final concentration of 0.2% (w/v), which equates to 13.3 mM. For activity determination of Hyd, the TGYEP medium, pH 6.5 was used, which includes 1% (w/v) tryptone, 0.5% (w/v) yeast extract, 0.1 M potassium buffer pH 6.5 and 0.8% (w/v) glucose (34). For experiments with different carbon sources M9-minimal medium was used. Concentrations of carbon sources were 0.4% (w/v) for cellobiose, 0.4% (w/v) for glycerol and 15 mM for fumarate, or as otherwise indicated. IPTG was added to a final concentration of 0.2 mM.

**Table 3:**
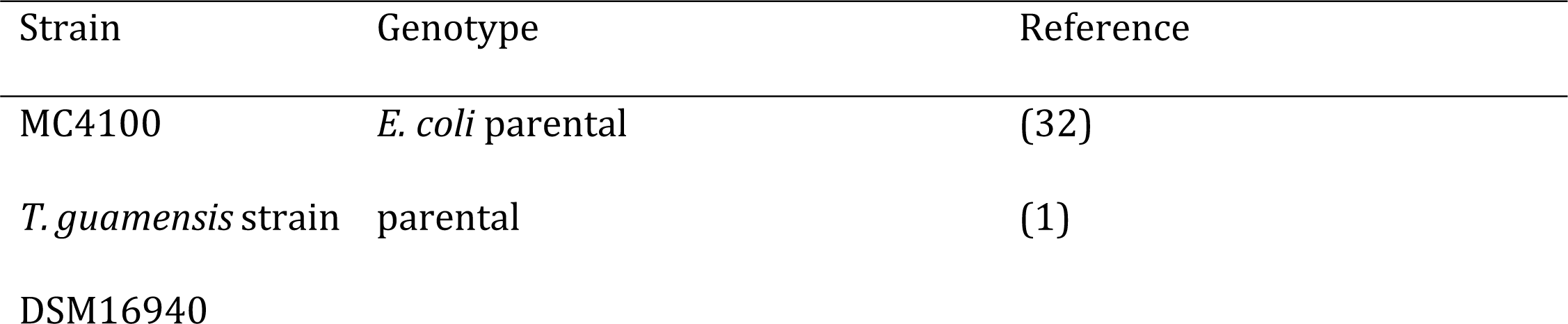
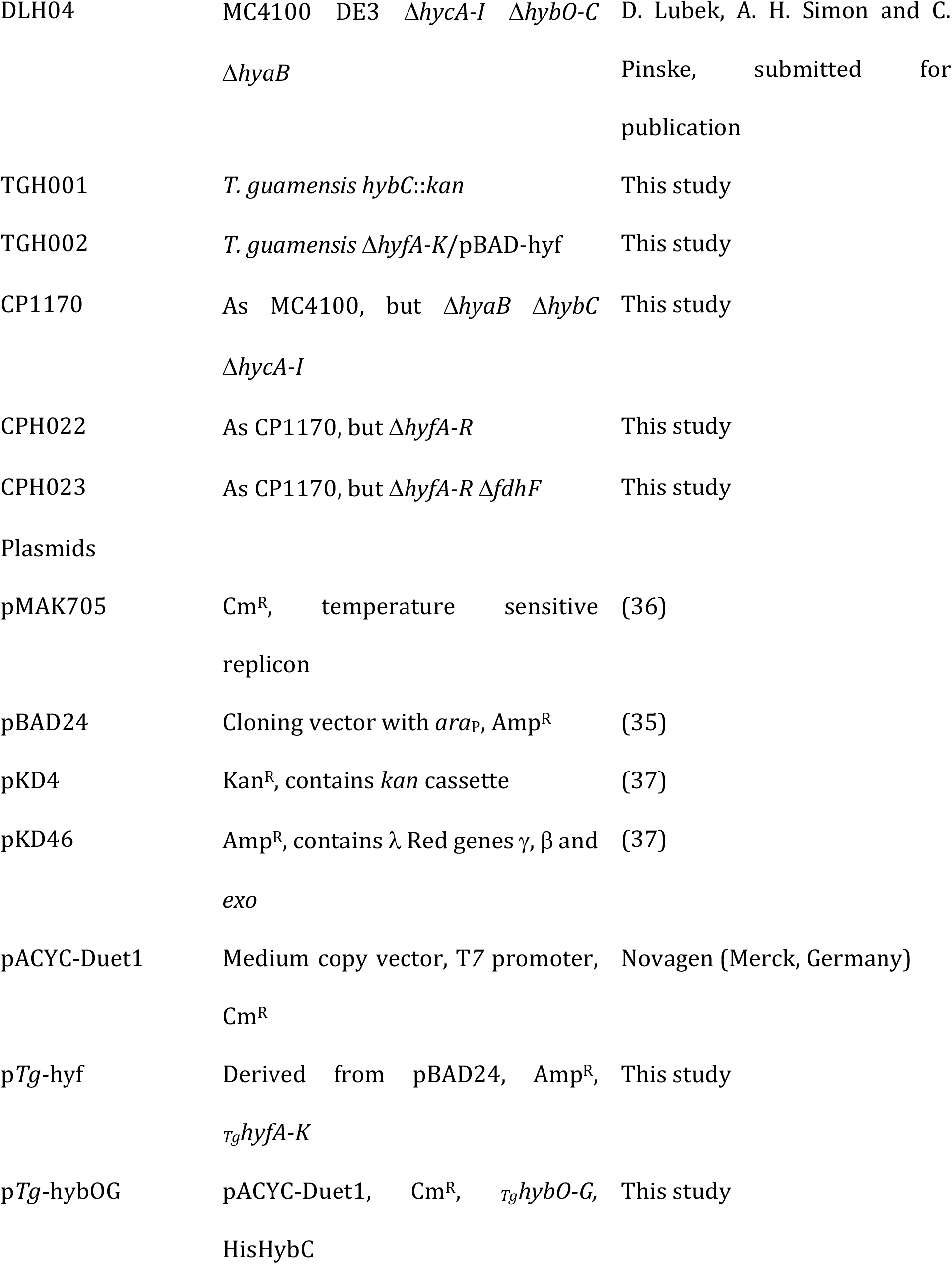
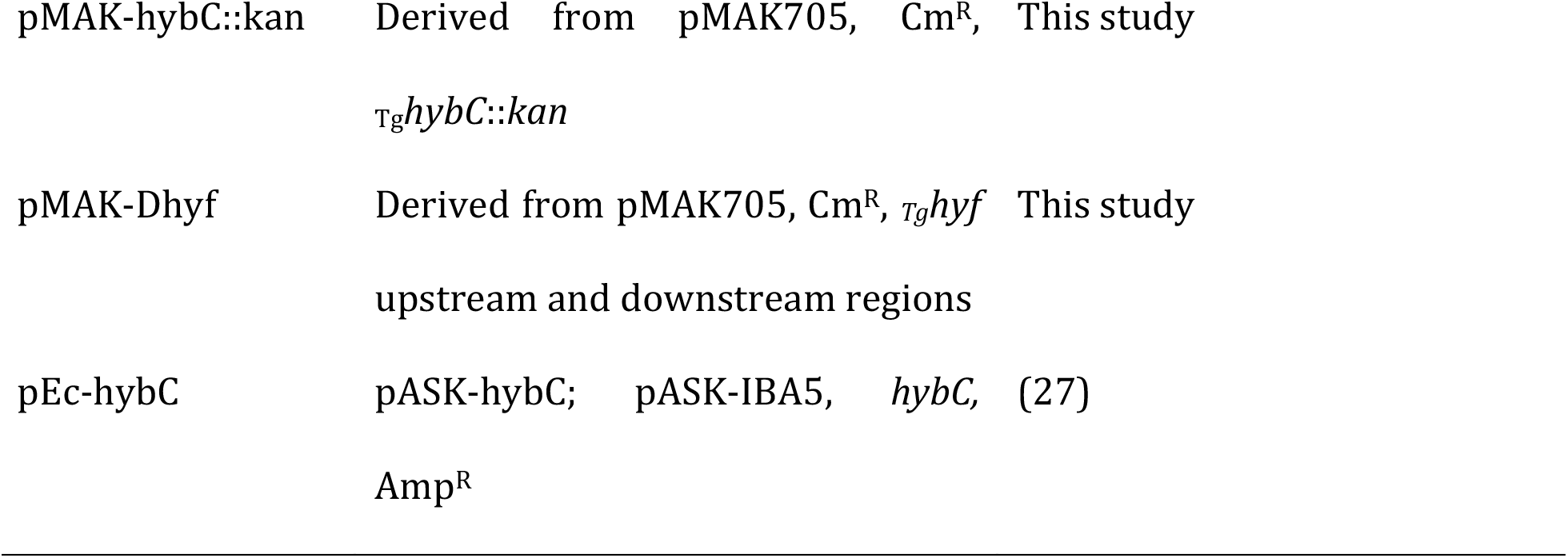
Strains and plasmids used in the study.

### Strain and plasmid construction

The *T. guamensis hyfA-K* operon comprising a total of 12 genes was cloned as three PCR fragments that were obtained from *T. guamensis* genomic DNA amplification and had the sizes 5058 bp, 3985 bp and 4048 bp. These were then assembled within PCR-amplified pBAD24 (35) with the NEBuilder kit, according to manufacturer’s instructions (NEB), which generated plasmid p*Tg-*hyf. The oligonucleotides used for this construction are listed as number 1-8 in Tab. 4.

**Table 4:**
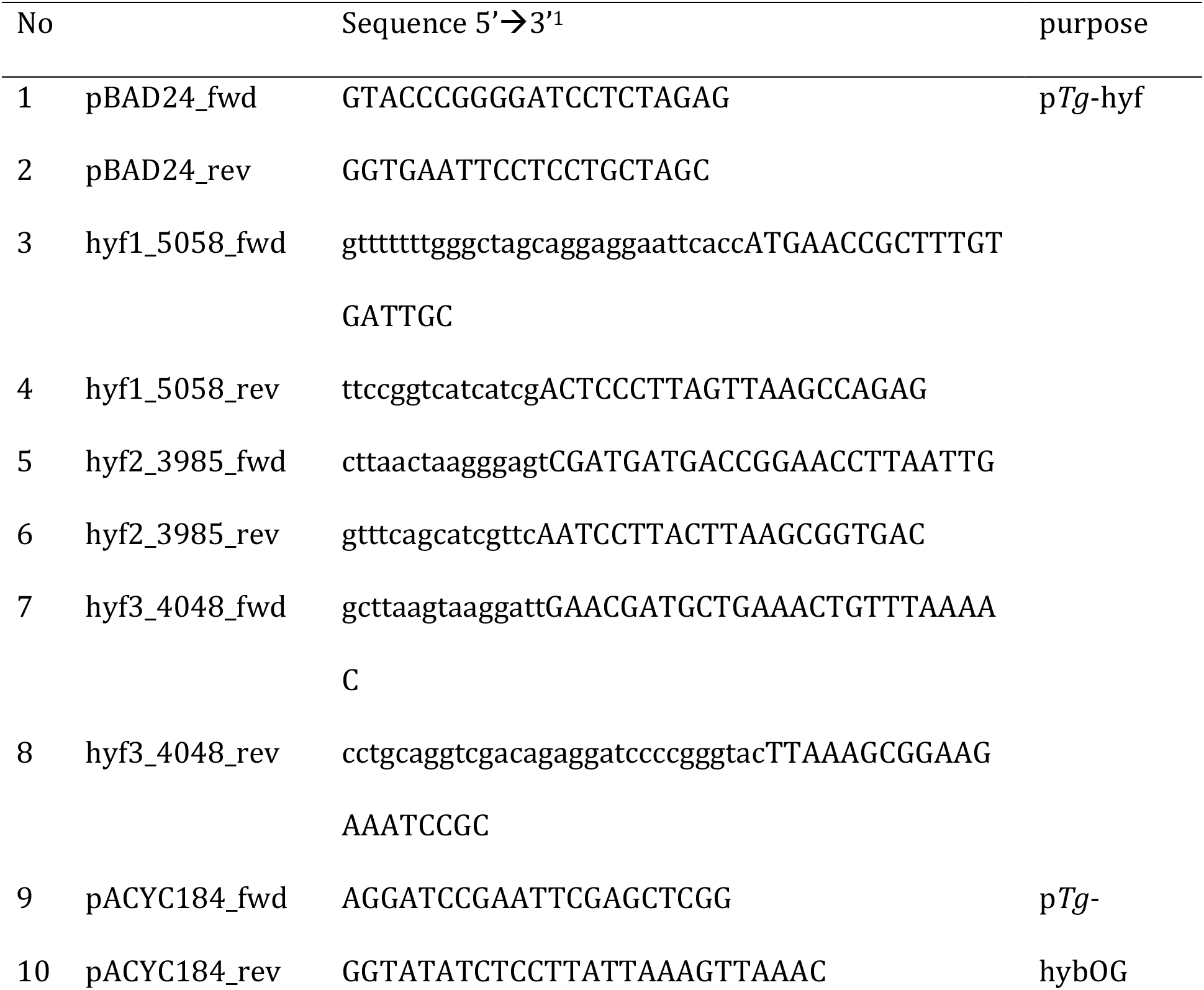
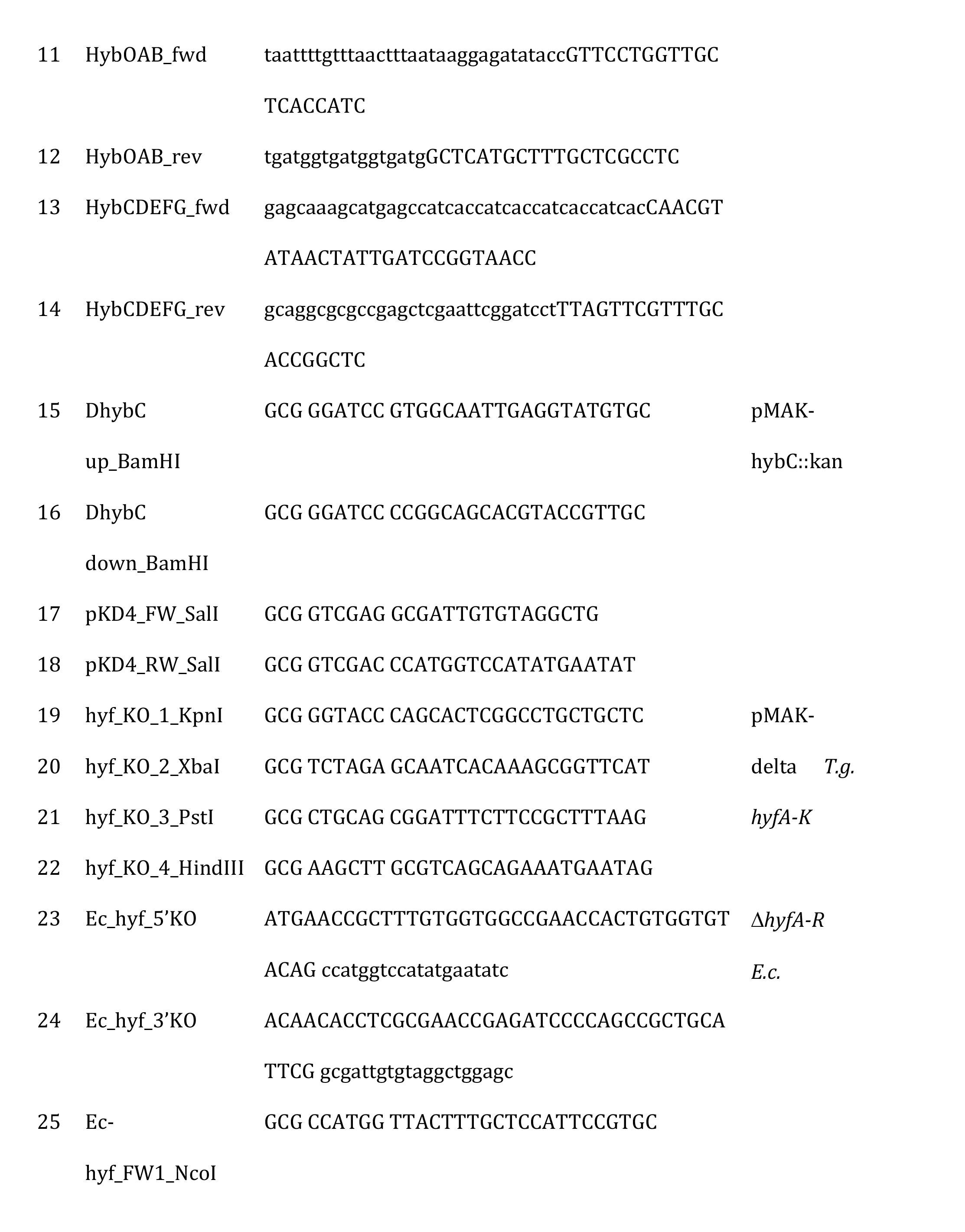
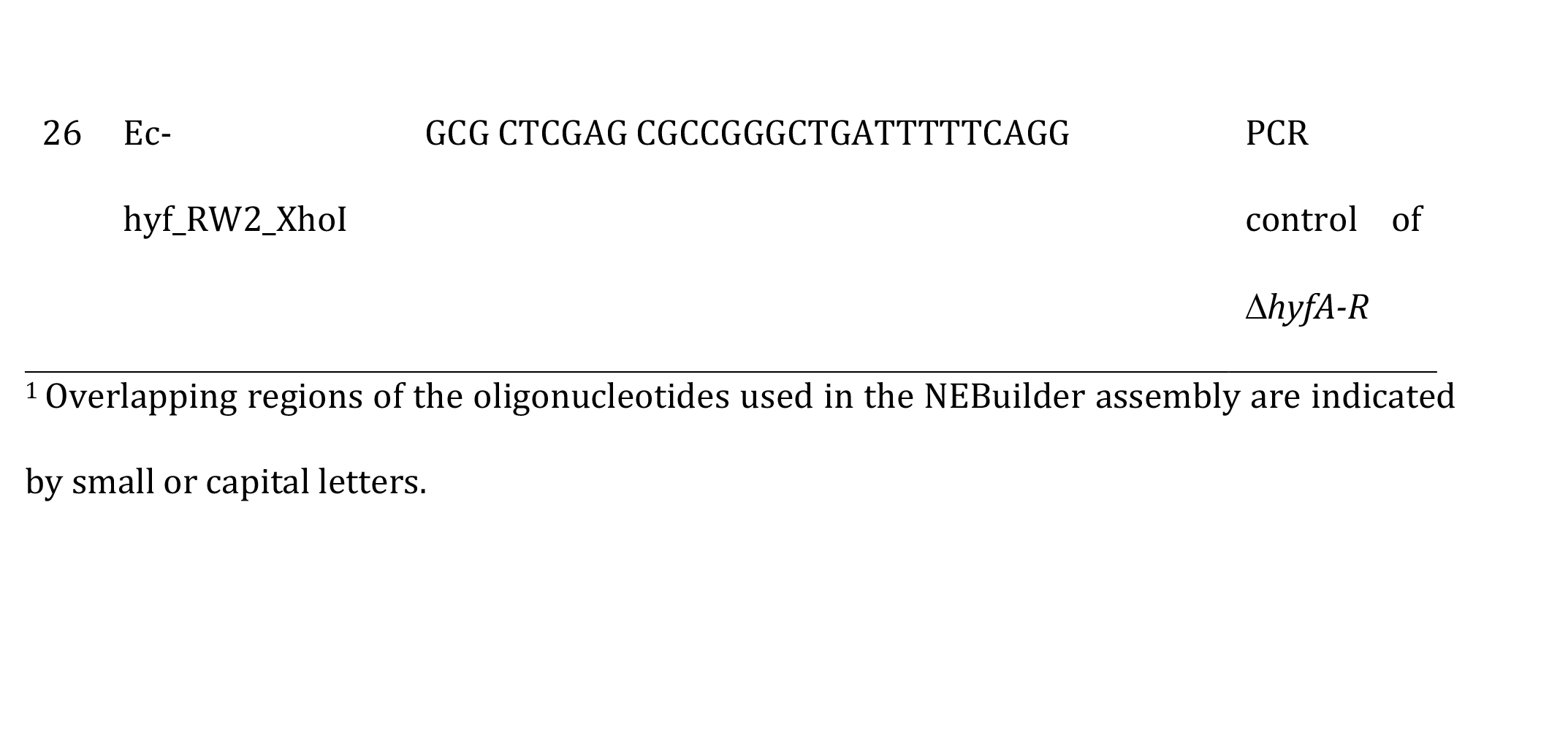
Oligonucleotides used in this study.

Similarly, the *T. guamensis hybO-G* operon including a 650 bp upstream region was amplified as two fragments with the oligonucleotides 11+12 and 13+14 (Tab. 4), which introduced a sequence coding for a His_8_-stretch *N*-terminally on HybC. The fragments were assembled on PCR-amplified (oligonucleotides 9+10) pACYC-Duet1 vector by the NEBuilder method (NEB) resulting in plasmid p*Tg*-hybOG.

Construction of *T. guamensis* mutants was done according to the pMAK705 method as described before (36). For the *hybC*∷*kan* insertion mutant, the *hybC* gene was amplified from chromosomal DNA using the oligonucleotides 15+16 (Tab. 4), cloned into pJET1.2 and the intrinsic SalI site then used to insert the kanamycin cassette from vector pKD4 (37), which had been previously amplified using the oligonucleotides 17+18 (Tab. 4). The construct was then moved as BamHI/BamHI fragment into pMAK705 and the resulting pMAK-hybC∷kan used for recombination by selecting for the kanamycin resistance during the final step. The resulting strain was named TGH001 (*T. guamensis hybC*∷*kan*).

To introduce the *hyfA-K* deletion into the *T. guamensis* genome, the upstream and downstream 500 bp regions of the operon were amplified and cloned successively into pMAK705 as KpnI/XbaI and PstI/HindIII fragments using the oligonucleotides 19+20 and 21+22 (Tab. 4), resulting in pMAK-Dhyf. The chromosomal *hyf* deletion in *T. guamensis* was then constructed in the presence of the p*Tg*-hyf plasmid and 0.2% (w/v) L-arabinose. The resulting strain was named TGH002 (*T. guamensis* Δ*hyfA-K*/pBAD-hyf).

Genomic mutants were checked by colony PCR.

The deletion of *hyfA-R* operon in *E. coli* left *focB* still intact. The strain CP1170 (Δ*hyaB* Δ*hybC* Δ*hycA-I*) was transformed with λ-red carrying plasmid pKD46, grown according to (37), and a PCR fragment (oligonucleotides 23+24 in Tab. 4), including the Cm^R^ resistance from pKD3 and 40 nucleotides flanking the deletion region. The authenticity of the resulting Cm^R^ clones was verified with oligonucleotides 25+26 (Tab. 4).

### Enzyme activities

Native PAGE was performed in 7.5% (w/v) polyacrylamide gels as described and stained for hydrogenase enzyme activity with 0.5 mM benzyl viologen and 1 mM 2,3,5-triphenyl tetrazolium chloride under a 2% (v/v) H_2_ atmosphere (24).

Spectrophotometric hydrogenase enzyme assays were performed as described (38) with crude extracts that were obtained after sonication (20W, 2 min, 0.5 sec pulses) or with whole cells. Cuvettes were filled with 800 μl 4 mM benzyl viologen in 50 mM MOPS buffer, pH 7.0, and the remaining gas phase flushed with H_2_. The reduction of benzyl viologen was monitored at 600 nm and an ε_M_ of 7400 M^-1^ cm^-1^ was assumed. One unit of activity represents the conversion of one μmol substrate per min.

H_2_ headspace content was determined with a Shimadzu-2010 gas chromatography system equipped with a Shin Carbon Micropacked column ST80/100 column as described (31). The amount of H_2_ was normalized to the OD_600 nm_ of the culture and the used culture volume. The continuous H_2_ production from whole cells was monitored using a Clark-type electrode (Hansatech instruments) modified to detect H_2_ and filled with 2 ml of degassed 50 mM MOPS buffer pH 7.0. The reaction was started by the addition of formate to a final concentration of 50 mM. Calibration was done with H_2_-saturated buffer according to (39). Protein concentration was determined according to the method of Lowry (40).

## Author contributions

UL and CP have performed the experiments, CP has conceptualized the experiments, and drafted the manuscript.

## Acknowledgments and funding information

The project was funded by the Deutsche Forschungsgemeinschaft project PI 1252/2 to CP. We wish to thank Gary Sawers for critical and helpful discussions.

